# Morphology of male genitalia and legs reveals the classification of Mesozoic Zoraptera (Insecta)

**DOI:** 10.1101/2025.01.08.631689

**Authors:** Petr Kočárek, Ivona Kočárková, Robin Kundrata

**Affiliations:** Department of Biology and Ecology, Faculty of Science, University of Ostrava, Chittussiho 10, CZ-710 00 Ostrava, Czech Republic; Department of Zoology, Faculty of Science, Palacky University, 17. listopadu 50, CZ-771 46 Olomouc, Czech Republic

**Keywords:** Fossils, amber, systematic paleontology, taxonomy, Cretaceous, Polyneoptera, Insects

## Abstract

Zoraptera is a small insect order with less than 50 described species from mostly tropical and subtropical regions of the World. The current system of extant Zoraptera is based on the results of molecular phylogeny combined with the morphology of male genitalia, and supplemented by the characters on the male abdomen and a number of metatibial spurs. However, fossil representatives of Zoraptera have not yet been classified into the modern system and most of them remained in the collecting genus *Zorotypus* Silvestri, 1913, because the genitalia were not observable or examined in detail. In this study, for the first time, we describe and critically evaluate the male genitalia and other principal diagnostic characters of all available Mesozoic Zoraptera. Our results lead to the first proposal of the generic classification of Mesozoic Zoraptera. We describe two new genera, *Cretozoros* gen. nov. and *Burmazoros* gen. nov., reinstate *Paleospinosus* Kaddumi, 2005, stat. restit. from synonymy with the subgenus *Octozoros* Engel, 2003 (in *Zorotypus*), and elevate *Octozoros* Engel, 2003 to a genus level. *Cretozoros* gen. nov., *Paleospinosus* stat. restit. and *Octozoros* stat. nov. are classified in Spiralizoridae: Latinozorinae, while *Burmazoros* gen. nov. and *Xenoburmiticus* Engel & Grimaldi, 2002 are classified in Zorotypidae: Zorotypinae. Altogether, it was possible to classify nine out of the 11 currently recognized species of Mesozoic Zoraptera. *Zorotypus hukawngi* Chen & Su 2019 is synonymized with *Cretozoros acanthothorax* (Engel & Grimaldi, 2002), comb. nov. and *Zorotypus hirsutus* Mashimo, 2018 is synonymized with *Octozoros robustus* (Liu, Zhang, Cai & Li, 2018), stat. restit., comb. nov., which is simultaneously restituted from the synonymy with *Octozoros cenomanianus* (Yin, Cai & Huang, 2018), comb. nov. The classification of Mesozoic Zoraptera into the modern system enables us to better understand the diversity of their internal lineages during the early evolution of this enigmatic insect order.

## Introduction

Zoraptera, or angel insects, represent one of the smallest insect orders, with only 47 extant and 16 extinct described species [1–4]. Zoraptera are small, soft-bodied, primarily winged polyneopteran insects. However, winged specimens are rare in the populations and most of collected specimens are wingless. The wing dimorphism is one of a few autapomorphies of the order, correlated with the presence or absence of compound eyes and ocelli, and the presence or absence of a distinct pigmentation, with alate specimens being distinctly darker [5]. Zoraptera are predominantly known from the subtropics and tropics, where they live mainly under the bark of rotting wood. Although zorapterans seem to be a common insect in the tropics, they are rarely collected due to their cryptic lifestyle and visual inconspicuousness [6].

Zoraptera represent an old evolutionary lineage with a Paleozoic origin [7–10]. Although Misof et al. [11] suggested that Zoraptera diverged from a common ancestor shared with Dermaptera in the Middle Jurassic (ca 180–160 Ma), later analyses [9] shifted that split to as early as 370 Ma. Most recently, Tihelka et al. [10] recovered Zoraptera as the earliest-diverging polyneopteran order and based on a set of justified fossil and stratigraphic calibrations, they proposed the Early to Late Devonian origin of Zoraptera (402–365 Ma). Therefore, more lines of evidence suggest that the origin of Zoraptera dates back to the Palaeozoic and that Zoraptera were already diversified and widely distributed in Gondwana at the time of the breakup of this supercontinent [1, 12].

Zorapteran uniformity in general morphology has led to the persistence of a conservative classification of extant Zoraptera, with only a single nominotypical genus *Zorotypus* Silvestri, 1913 in a single family Zorotypidae for more than a century [5]. Kočárek et al. [13] and Matsumura et al. [8] conducted molecular phylogenetic studies using a combination of nuclear and mitochondrial markers. Both of these independent analyses revealed two major phylogenetic lineages, which Kočárek et al. [13] classified as families Zorotypidae and Spiralizoridae, each of them further subdivided into two robustly supported subfamilies [13]. The updated classification was supported by synapomorphies in the structure and shape of the male genitalia and other taxonomically valuable characteristics including the morphology of the apex of the male abdomen and a number of spurs on metatibia. Unfortunately, molecular-genetic methods cannot be applied to insect fossils trapped in amber [14, 15]; therefore, their systematic placement must be inferred from the morphological characters. In Zoraptera, the most important characters are found in the male genitalia, but these are usually enclosed in the abdomen and are only rarely visible [16].

Altogether, 12 fossil species of Zoraptera are currently recognized from the Mesozoic [1]. One species is reported from the lower Cretaceous Jordanian amber (Albian) [17], and 11 species are known from the upper Cretaceous amber (Cenomanian) of northern Myanmar [1]. Fossil zorapterans are classified partly in *Zorotypus sensu stricto*, partly in a monotypic genus *Xenozorotypus* Engel & Grimaldi, 2002, and partly in the exclusively fossil subgenus *Octozoros* Engel, 2003, which was erected for species with eight antennomeres [5, 18]. All of these taxa are included in a single family, Zorotypidae, although it is evident from their morphology that they belong to different phylogenetic lineages and should be reclassified [8, 13].

In this study, for the first time, we critically evaluate the principal diagnostic characters of the Mesozoic Zoraptera, including male genitalia. Our conclusions are based partly on the available literature and images of Mesozoic species, and partly on a new set of fossils in Burmese amber in which the diagnostically significant parts of the genitalia were observable. Based on detailed examination of male genitalia and other important characters, we were able to classify most Mesozoic Zoraptera species into the currently accepted system, which was created based on extant taxa. The relationships of extinct and recent zorapteran groups are evaluated and discussed.

## Results

### Systematic paleontology

Order Zoraptera Silvestri, 1913

Family Zorotypidae Silvestri, 1913

Subfamily Zorotypinae Silvestri, 1913

#### Genus *Burmazoros* gen. nov

urn:lsid:zoobank.org:act:089A6025-92CB-4242-97CE-C0FFB7980D85

##### Type species

*Zorotypus denticulatus* Yin, Cai & Huang, 2018, here designated.

##### Etymology

The generic name refers to the occurrence of this taxon in Burmese amber, in combination with a suffix derived from the word base of the order name. Gender masculine.

##### Diagnosis

*Burmazoros* gen. nov. is characterized by the following unique combination of characters: antennae 9-segmented; metafemur with middle spine larger than basal spine; metatibia with three acute spines at middle, and at preapical and apical portions; ctenidia on T10 missing, T10 and T11 each bear a small conical median projection (MP) slightly curved upwards. Male genitalia asymmetrical, rod-shaped accessory sclerites developed laterally. *Burmazoros* gen. nov. differs from recent *Zorotypus* Silvestri, 1913 in reduced wing venation: CuA missing in fore wings; hind wings with missing M1+2, and short M fused with R in the middle; R not reaching wing margin.

##### Systematic placement

*Burmazoros* gen. nov. is herein classified in Zorotypidae: Zorotypinae based on the following characters: asymmetrical male genitalia, not developed intromittent organ, and three metatibial spurs. We suppose that this genus is related to recent genera *Zorotypus* and *Usazoros* Kukalova-Peck & Peck, 1993.

##### Species included

*Burmazoros denticulatus* Yin, Cai & Huang, 2018 (=*Zorotypus oligophleps* Liu, Zhang, Cai & Li, 2018).

##### Distribution

Myanmar, Kachin State, Myitkyina District, Hukawng Valley, Burmese amber (Upper Cretaceous, lower Cenomanian).

#### *Burmazoros denticulatus* (Yin, Cai & Huang, 2018), comb. nov

*Zorotypus denticulatus* Yin, Cai & Huang, 2018: 169.

=*Zorotypus oligophleps* Liu, Zhang, Cai & Li, 2018: 260.

##### Comments

This species is known only from the type specimen, which is a well-preserved alate male [19]. The metafemur has a unique arrangement of spines; metatibia bears three strong spurs in the distal third. Male genitalia are partly visible, asymmetrical, rod-shaped accessory sclerites are developed laterally. T10 and T11 are with short mating hooks.

Yin et al. [20] synonymized *Zorotypus oligophleps* Liu, Zhang, Cai & Li, 2018 with *Zorotypus denticulatus* Yin, Cai & Huang, 2018. These species share a similar form of the head, the pronotum, identical configuration and proportions of the antennomeres, and a similarly reduced wing venation [19]. The only difference is in the number of metatibial spurs, *Burmazoros denticulatus* has three spurs, and Z. *oligophleps* has two spurs [20]. Although the number and arrangement of the metatibial spur is an important diagnostic character, the middle spur in the holotype of *Z. oligophleps* may actually be actually broken off. Until the number of spurs on the newly found individuals of this species is confirmed, we provisionally keep Z. *oligophleps* in synonymy.

#### Genus *Xenozorotypus* Engel & Grimaldi, 2002

*Xenozorotypus* Engel & Grimaldi, 2002: 12.

##### Type species

*Xenozorotypus burmiticus* Engel & Grimaldi, 2002, by original designation.

##### Systematic placement

*Xenozorotypus* is herein classified to Zorotypidae: Zorotypinae. *Xenozorotypus* shares three metatibial spurs, however, male genitalia are not observable in the only known specimen of this species [21]. We suppose it is related to the recent genera *Zorotypus* and *Usazoros*.

##### Species included

*Xenozorotypus burmiticus* Engel & Grimaldi, 2002

##### Distribution

Myanmar, Kachin State, Myitkyina District, Hukawng Valley, Burmese amber (Upper Cretaceous, lower Cenomanian).

#### Xenozorotypus burmiticus Engel & Grimaldi, 2002

*Xenozorotypus burmiticus* Engel & Grimaldi, 2002: 12.

##### Comments

This species is known only from the holotype, which is not a well-preserved male specimen [21]. The metafemur shows an exceptionally deep ventral furrow that extends from the apex to the midpoint; metatibia has three strong spurs regularly spaced in the distal two-thirds, and a basal spine is present on the ventral surface of metatibia. T10 bears a procurved mating hook. Specific is the venation of the hind wing with M3+4 present. Genitalia are not observable.

Family Spiralizoridae Kočárek, Horká & Kundrata, 2020

Subfamily Latinozorinae Kočárek, Horká & Kundrata, 2020

#### Genus *Octozoros* Engel, 2003, stat. nov

(Figs 1,2)

**Figure 1.**
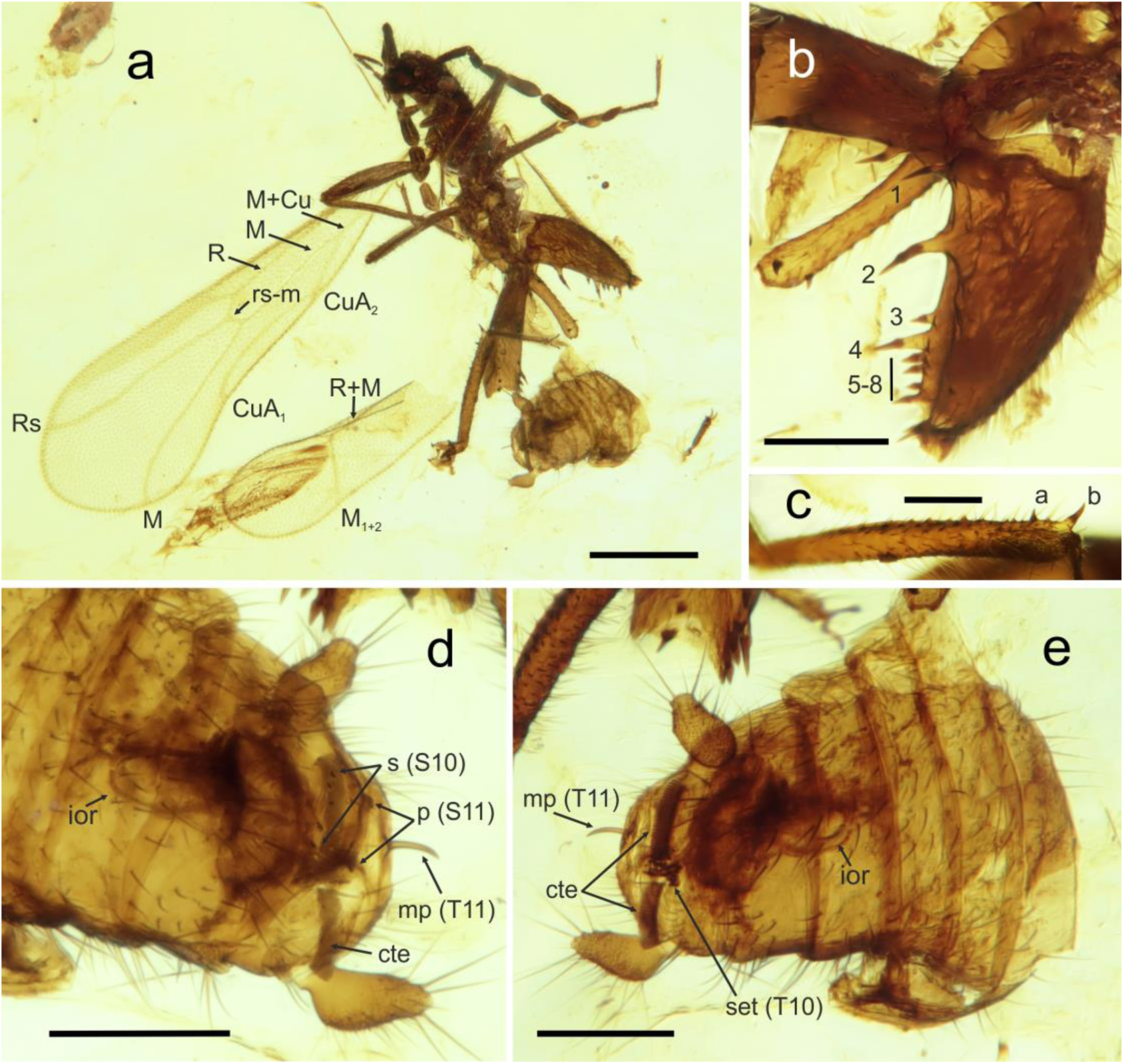
Habitus and details of *Octozoros cenomanianus* (Yin, Cai & Huang, 2018), comb. nov., specimen PK344Bu, male. (a) General habitus. (b) Detail of left metafemur, lateral view. (c) Detail of right metatibia, lateral view. (d) Abdomen, ventral view. (e) Abdomen, dorsal view. Abbreviations: a, b, metatibial spurs; cte, ctenidium; Cu, cubitus vein; CuA, anterior cubitus vein; ior, intromittent organ; M, media vein; R (Rs), radius vein; mp, median up-curved projection; p, projections; s, spines; set, thickened setae; S10–S11, sternites; T10– T11, tergites; 1–8, metafemoral spurs. Scale bars: 0.5 mm in (a), 0.2 mm in (b, d, e), 0.1 mm in (c).

**Figure 2.**
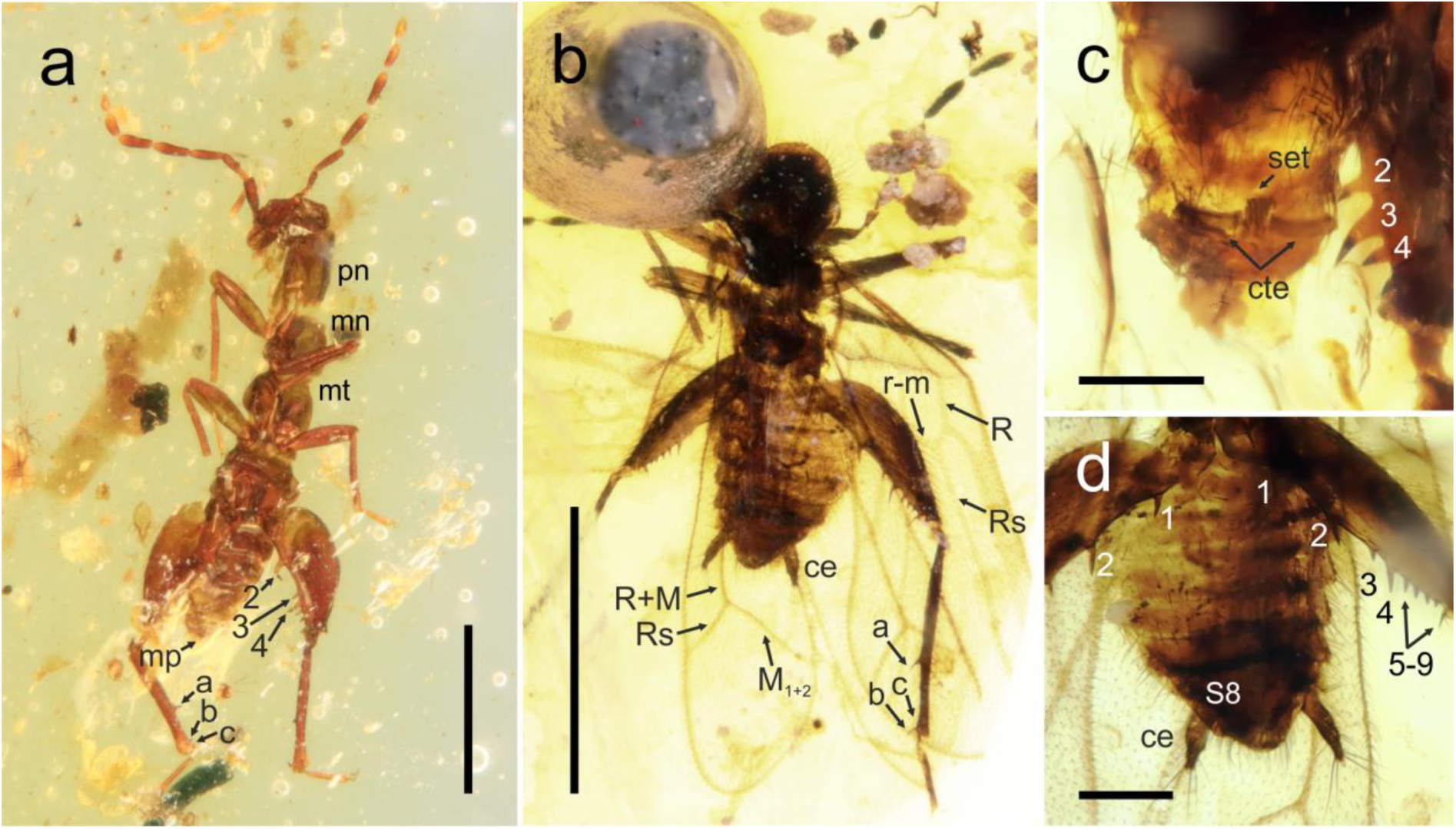
Habitus and details of *Octozoros robustus* (Liu, Zhang, Cai & Li, 2018), stat. restit., comb. nov., (a, c) specimen PK343Bu, male, (b, d) specimen PK348Bu, female. (a) General habitus of apterous male, dorsal view. (b) General habitus of winged female, ventral view. (c) Detail of male abdomen, ventral view. (d) Detail of female abdomen, ventral view. Abbreviations: a, b, c, metatibial spurs; cte, ctenidium; M, media vein; mn, mesonotum; mp, median up-curved projection; mt, metanotum; pn, pronotum; R (Rs), radius vein; set, tickened setae; S8, sternite 8; 1–9, metafemoral spurs. Scale bars: 1.0 mm in (a, b), 0.2 mm in (c, d).

*Octozoros* Engel, 2003: 148 (as subgenus of *Zorotypus*).

##### Type species

*Zorotypus nascimbenei* Engel & Grimaldi, 2002, by original designation (in the subgenus *Octozoros* Engel, 2003).

##### Updated diagnosis

*Octozoros* is characterized by the following unique combination of characters: antennae 8-segmented; forewings with fully developed venation, R divided into R and Rs in midpart, Rs continuing from the radial stem and connected with M by short Rs-M crossvein, M and Rs reaching posterior wing margin near the wing apex, CuA_1_ present; ventral margin of metafemur not furrowed or with shallow furrow; hind tibiae with two robust spurs, one in distal third to fifth, second distally; in some species, e.g. *O. robustus* (Liu, Zhang, Cai & Li, 2018) and *O. pecten* (Mashimo, Mueller & Beutel, 2019), another tiny apical spine developed inside the ventral surface; in males, T10 with two rows of thick setae arranged as a comb (ctenidium) on both sides, the central region of T11 with long and thin upcurved MPs, which are missing on T10; male genitalia symmetrical with robust basal part and tongue-like anterior process encircled by intromittent organ, lateral rod-shaped accessory sclerites missing.

Among the Latinozorinae, *Octozoros* differs from *Cretozoros* gen. nov. in the presence of CuA_1_ on the fore wings, ctenidia on T10, and only T11 with upcurved MP. *Octozoros* differs from recent *Latinozoros* Kukalova-Peck & Peck, 1993 in the number of MPs in the male abdomen (*Octozoros* has MP only on T11, while *Latinozoros* has MPs on both T10 and T11) and in the number of antennomeres (antennae of *Octozoros* are composed of 8 antennomeres in adults of both sexes, while antennae of *Latinozoros* are always composed of 9 antennomeres in adults).

##### Systematic placement

Subgenus *Octozoros* (in *Zorotypus*) is herein elevated to a genus level and classified in Spiralizoridae: Latinozorinae. *Octozoros* shares with *Latinozoros* a similar pattern of symmetrical male genitalia with the basal plate encircled by the intermittent organ, as well as fore wings with developed CuA_1_ vein. Therefore, we consider this taxon sister to the *Cretozoros*/*Latinozoros* clade (Fig. 4).

##### Species included

*Octozoros cenomanianus* (Yin, Cai & Huang, 2018); *O*. *nascimbenei* (Engel, Grimaldi 2002); *O. pecten* (Mashimo, Mueller & Beutel, 2019); *O. robustus* (Liu, Zhang, Cai & Li, 2018), stat. restit. (=*Z. hirsutus* Mashimo, 2018, syn. nov.).

##### Distribution

Myanmar, Kachin State, Myitkyina District, Hukawng Valley, Burmese amber (Upper Cretaceous, lower Cenomanian).

#### *Octozoros cenomanianus* (Yin, Cai & Huang, 2018), comb. nov

(Fig. 1)

*Zorotypus cenomanianus* Yin, Cai & Huang, 2018: 169.

##### Material examined

One adult specimen: alate male, PK344Bu (Burmese amber).

##### Supplemented description

Based on the study of new material, we supplement the description of *C. cenomanianus* with the following morphological characters, which were not observable in the holotype [19]. Both specimens known so far are adult males; immature stages and the female specimens remain undescribed.

##### Legs

Metatibia with two stout spines, one in the apical fifth, the second distally (Fig. 1a–c). The spines are not orientated to each other on the axis of the tibia, but the proximal spine is located on the outer edge of the tibia and the distal spine on the inner edge (Fig. 1c). The spines are therefore orientated obliquely to each other (Fig. 1b). Metapretarsus with pair of thin bristles; empodium reduced to a short hair-like structure.

##### Male genitalia

Symmetrical, composed of robust basal part without recognized details, and proximally oriented protrusion (basal plate) horizontally encircled by an intromittent organ.

#### *Octozoros nascimbenei* (Engel & Grimaldi, 2002), comb. nov.

*Zorotypus nascimbenei* Engel & Grimaldi, 2002: 7.

##### Comments

This species is known only from the type specimen, which is an alate female [21]. Species-specific is the pattern of the metafemur spination, with spines 1 and 2 robust, and the remaining spines short, reaching maximally half of the length of spines 1 and 2. The metatibia bears two strong spurs in the distal third. The pronotum has a shallow depression in the apical margin. Males are not recorded, therefore the characters on male genitalia, arrangement of ctenidium on T10, and mating hooks on T10 and T11 (presence and shape) cannot be evaluated. The species is classified in *Octozoros* stat. nov. by the combination of the following observable characters: 8-segmented antennae, CuA_1_ present, and hind tibiae with two spurs in distal third.

#### *Octozoros robustus* (Liu, Zhang, Cai & Li, 2018), stat. restit., comb. nov

*Zorotypus robustus* Liu, Zhang, Cai & Li, 2018: 260.

=*Zorotypus hirsutus* Mashimo, 2018: 563, syn. nov.

(Fig. 2)

##### Material examined

Four adult specimens: two alate females, PK348Bu, PK164Bu, one apterous male, PK343Bu, and one apterous female, PK336Bu (all Burmese amber).

##### Taxonomic remarks

Yin et al. [20] synonymized *Zorotypus robustus* with *Zorotypus cenomanianus* (currently in the genus *Octozoros*). Based on the comparison of original descriptions as well as on the study of new material of both species (see Material and Methods), these two species were recognized as distinct. Therefore, *Z. robustus* is resurrected from synonymy and is considered to be a valid species in the genus *Octozoros*. Both species have very similar patterns of spurs arrangement on metafemur, but differ in the arrangement of spurs on metatibiae (compare Fig. 1a,c and Figures 1a,c in Liu et al. [22]). Both species also differ in the shape of the pronotum, which is quadrate in *O. robustus* but distinctly longer than wide in *O. cenomanianus* (compare Figures 1a,c in Liu et al. [22] and 1a,b,c in Yin et al. [19]). The difference is also in the antennomeres 4–6, which are slender in *O. robustus* (Fig. 2a,b) but more robust in *O. cenomanianus* (Fig. 1a), and in cerci, which are slender in *O. robustus* (Fig. 2b,d) but more robust in *O. cenomanianus* (Fig. 1d,e).

*Zorotypus hirsutus* is here synonymized with *Octozoros robustus* based on detailed morphological comparisons between published descriptions and available fossil material (see above and Material and Methods) which led to the finding that all diagnostic characters match and there are actually no morphological differences between these species. Both species were described in the same year and neither author mentioned the existence of the other of these species in the original publications, and it is obvious that the authors were unaware of the concurrent description.

#### *Octozoros pecten* (Mashimo, Mueller & Beutel, 2019), comb. nov

*Zorotypus (Octozoros) pecten* Mashimo, Mueller & Beutel, 2019: 566.

##### Comments

This species is known only from the type specimen, which is a well preserved alate male [22]. Genus-level diagnostic characters are well observable, including the 8-segmented antennae, CuA_1_ present on fore wings, hind tibiae with two spurs in the distal third, and the additional tiny apical spine developed on the inside of the ventral surface, ctenidia present on both sides of T10, and the mating hook present on T11. Genitalia not observable. *Octozoros pecten* is similar to *O. cenomanianus*, but it differs by the presence of a group of thick setae in the middle of T10.

#### Genus *Cretozoros* gen. nov

urn:lsid:zoobank.org:act:2FC09AE9-C2E5-4898-8DB7-0B30333123FE

(Fig. 3)

**Figure 3.**
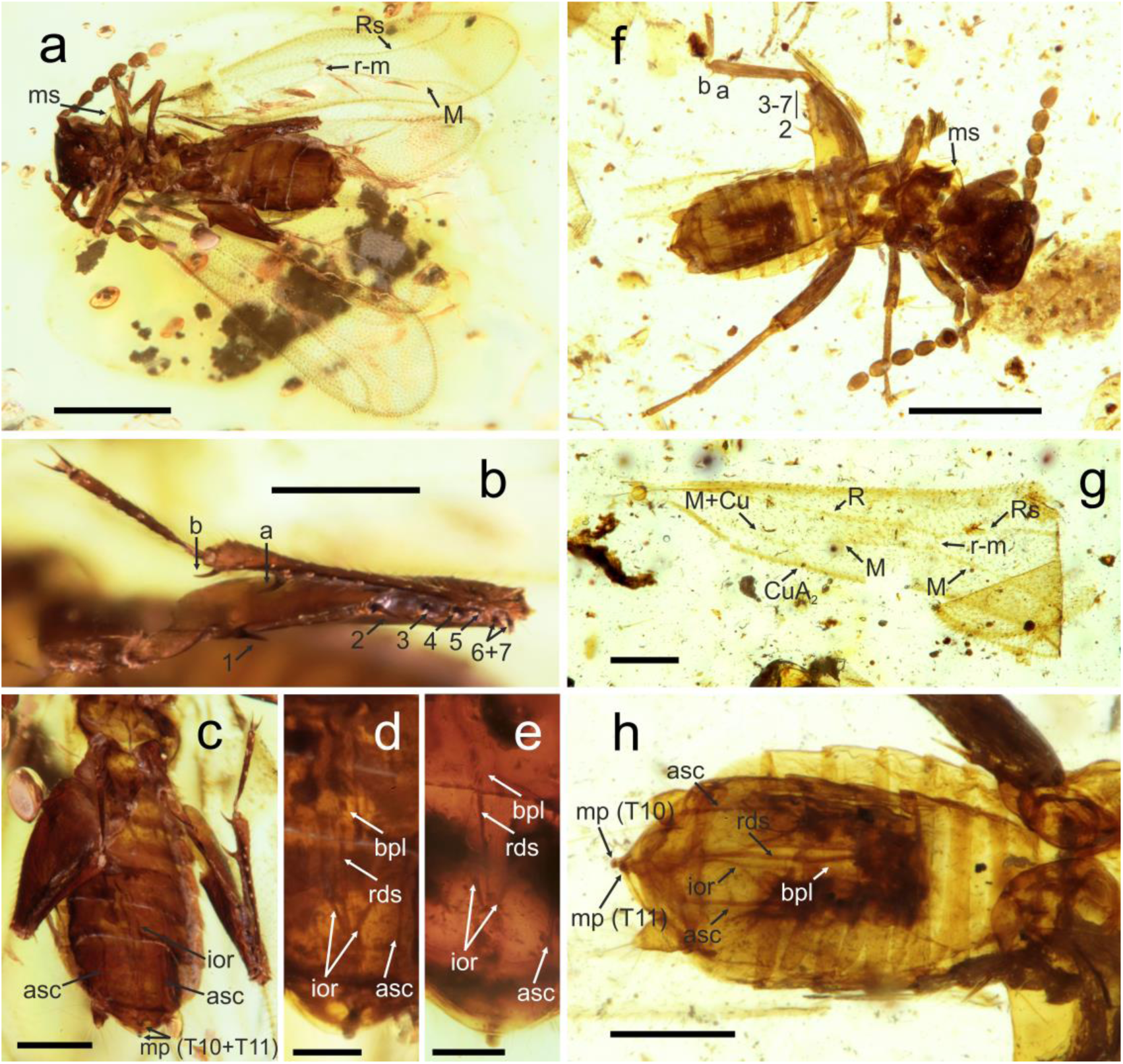
Habitus and details of *Cretozoros acanthothorax* (Engel & Grimaldi, 2002), comb. nov. (a–e) specimen PK342Bu, male; (e–h) specimen PK347Bu, male. (a, f) General habitus. (b) Detail of right metafemur and metatibia, ventral view (c, h) Abdomen, ventral view. (d, e) Detail of male genitalia, with different focus layers and lighting, ventral view. (g) Left forewing, ventral view. Abbreviations: a, b, metatibial spurs; asc, lateral accessory sclerites; bpl, basal plate; Cu, cubitus vein; CuA, anterior cubitus vein; ior, intromittent organ; M, media vein; ms, mesonotal spine; R (Rs), radius vein; mp, median projection; rds, median rod-like sclerites; S10–S11, sternites; T10–T11, tergites; 1–7, metafemoral spurs. Scale bars: 0.5 mm in (a, f), 0.2 mm in (b, c, g, h), 0.1 mm (d, e).

##### Type species

*Zorotypus acanthothorax* Engel & Grimaldi, 2002, here designated.

##### Etymology

The generic name refers to an exclusive occurrence in the Cretaceous period, in combination with a suffix derived from the word base of the order name. Gender masculine.

##### Diagnosis

*Cretozoros* gen. nov. is characterized by the following unique combination of characters: antennae 8-segmented; forewings with reduced venation, R not developed, Rs continuing from the radial stem and not divided into R and Rs distally from rs-m crossvein and continuing as Rs, M and Rs reaching posterior wing margin near wing apex, CuA_1_ absent; ventral margin of metafemur without furrow; hind tibiae with two spurs in distal third; in males, ctenidia on T10 missing, central regions of T10 and T11 with short upcurved MPs (MP T11 not visible or missing in *C. pussilus* (Chen & Su, 2019)); male genitalia symmetrical with tongue-like anterior process encircled by intromittent organ, laterally developed rod-shaped accessory sclerites.

*Cretozoros* gen. nov. differs from *Octozoros* in the absence of CuA_1_ on fore wings, missing ctenidia on T10, and MPs on both T10 and T11. *Cretozoros* gen. nov. differs from recent *Latinozoros* in the presence of rod-shaped accessory sclerites laterally of male genitalia. The antennae of *Cretozoros* are composed of eight antennomeres in adults of both sexes, whereas the antennae of *Latinozoros* adults are always composed of nine antennomeres in adults.

##### Systematic placement

Genus *Cretozoros* gen. nov. is herein classified in Spiralizoridae: Latinozorinae. It shares the venation pattern of the wing (missing CuA1), the development of two spurs on the metatibiae, and the morphology of the male genital (developed basal plate encircled by intromittent organ) with recent *Latinozoros*, and we suppose they constitute a monophyletic group (Fig. 4).

**Figure 4.**
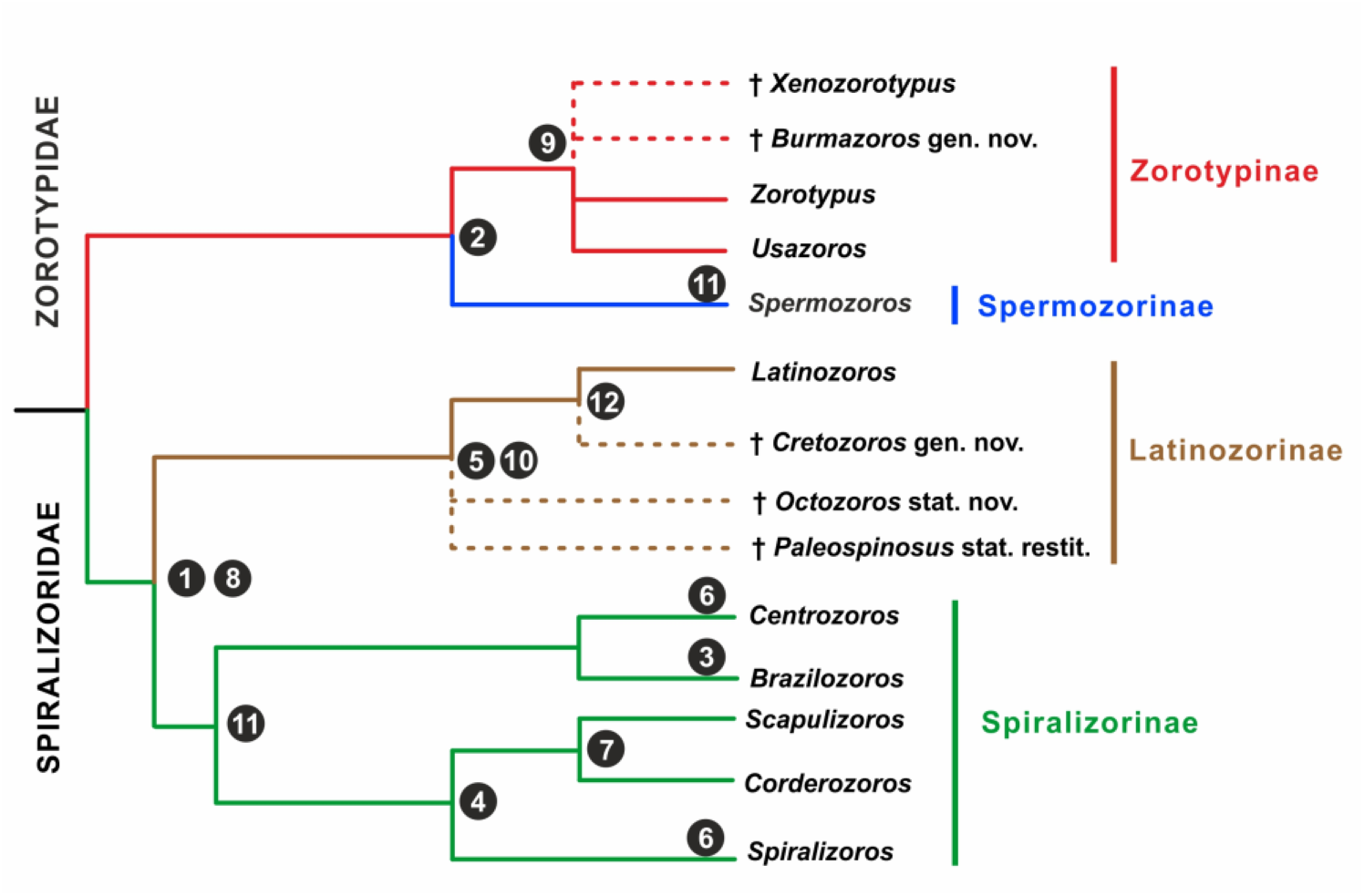
Simplified phylogenetic scheme of Zoraptera with probable placement of fossil genera *Cretozoros* gen. nov., *Burmazoros* gen. nov., *Octozoros* Engel, 2003, stat. nov., *Paleospinosus* Kaddumi, 2005, stat. restit., and *Xenozorotypus* Engel & Grimaldi, 2002. The phylogenetic tree is modified from Kočárek et al. [13]. Numbers refer to the following diagnostic characters: 1 – symmetrical male genitalia, 2 – asymmetrical male genitalia, 3 – intromittent organ absent; 4 – coiled or looped intromittent organ, 5 –intromittent organ horizontally looped, 6 – intromittent organ spirally coiled, 7 – intromittent organ dorsoventrally looped, 8 – basal plate of male genitalia developed, 9 – hind tibia with three spurs, 10 – hind tibia with two spurs, 11 – hind tibia without spurs, 12 – fore wings with CuA_1_ absent.

##### Species included

*Cretozoros acanthothorax* Engel & Grimaldi 2002 (= *Z. hukawngi* Chen & Su, 2019 [23], syn. nov.); *C. pusillus* Chen & Su, 2019 [16].

##### Distribution

Myanmar, Kachin State, Myitkyina District, Hukawng Valley, Burmese amber (Upper Cretaceous, lower Cenomanian).

#### *Cretozoros acanthothorax* (Engel & Grimaldi, 2002), comb. nov

*Zorotypus acanthothorax* Engel & Grimaldi, 2002: 10.

=*Zorotypus (Octozoros) hukawngi* Chen & Su 2019: 264, syn. nov.

(Fig. 3)

##### Material examined

Four adult specimens: three alate males, PK342Bu, PK347Bu, PK340Bu, and one alate female, PK346Bu (all Burmese amber).

##### Supplemented description

Based on the study of new material (see above, and Material and Methods below) and detailed morphological comparisons we found that there are no morphological differences between *Cretozoros acanthothorax* and *Zorotypus hukawngi*. Therefore, we synonymize *Z. hukawngi* with *C. acanthothorax*.

We supplement the description of *C*. *acanthothorax* with the following morphological characters, which were not observable neither in the holotype of *C. acanthothorax* nor in the holotype of *Z. hukawngi*. *Cretozoros acanthothorax* was described based on the male, and *Z. hukawngi was described* based on the female. The fossil material includes two males (PK347Bu, PK342Bu) and one female (PK346Bu).

##### Wing venation

Wings hyaline with dense pubescence, forewing length 1.64 mm, hind wing length 1.36 (Fig. 3a,g). The wing venation faint with most veins represented by fuscous lines, membrane hyaline with scattered minute setae except infuscation forming slightly sclerotized pterostigma in forewing; marginal setae on both fore and hind wings numerous and short, longer than setae on membrane; posterior margin of forewing with jugate setae in middle third. Forewings with reduced venation; R continuing from radial stem and not divided into R and Rs distally from rs-m crossvein (in the midpoint) and continuing as Rs; M and Rs reaching posterior wing margin near wing apex, CuA_1_ absent. Hindwing with M+R in anterior half, both R and M reaching wing margins, basal third of hind wing is not visible in any studied specimen; Cu absent.

##### Male abdomen

T10 smooth, without ctenidium, central regions of T10 and T11 distally with short peg-like MPs. Comment: the median projections are short and thus not well visible in some specimens – compare Fig. 3c,d,e,h. In the original description of *C*. *acanthothorax*, only MP T11 is depicted in Fig. 10 [21], but this fact is not mentioned in the description itself.

##### Male genitalia

Genitalia are symmetrical with a minute basal part and long tongue-like anterior process (basal plate) encircled by intromittent organ, axis of the basal plate with median rod-like sclerites, accessory rod-like sclerites are developed laterally (Fig. 3c,d,e,h).

#### *Cretozoros pusillus* (Chen & Su, 2019), comb. nov

*Zorotypus (Octozoros) pusillus* Chen & Su, 2019: 556.

##### Comments

This species is known only from the type series, which is composed of an alate male (holotype) and an alate female (paratype) embedded in the position of the copula in one amber piece [16]. Genus-level diagnostic characters are partly observable, including the 8-segmented antennae, and hind tibiae with two spurs developed in the distal third; the ctenidia are not developed. The mating hook is visible on T10, but the presence/absence of the mating hook on T11 cannot be evaluated due to the position of the fossil. The male genital is in an everted position and it is symmetrical; hovewer, the morphological details are not recognizable. Lateral rod-shaped accessory sclerites are visible. Species-specific is the arrangement of spines on the ventral surface of the metafemur.

#### Genus *Paleospinosus* Kaddumi, 2005, stat. restit

*Palaeospinosus* Kaddumi, 2007: 218.

##### Type species

*Paleospinosus hudae* Kaddumi, 2005, by original designation.

##### Updated diagnosis

*Paleospinosus* is characterized by the following unique antennae 8-segmented; anterolateral combination of characters: spines on the mesonotum missing; ventral margin of metafemur with a deep furrow that extends from apex to the middle; hind tibiae with two spurs in distal third; in males, T10 with two clumps of thick setae on both sides of distal margin; one long and thin upcurved MP developed, but this is not clear if this is the projection of T10 or T11; distal edge of S8 broadly emarginate with a peg-like projection in the middle. Wing venation is not distinctly recognizable in the holotype of type species, therefore cannot be used for the diagnosis.

Among Latinozorinae, males of *Paleospinosus* differ from those of other genera as follows: from *Cretozoros* gen. nov. in the presence of two clumps of thick setae on both sides of T10 distal margin, one long upcurved MP developed, and the distal edge of S8 broadly emarginate; from *Octozoros* in the missing ctenidia on T10 and broadly emarginate distal edge of S8; from recent *Latinozoros* in the number of MPs in the male abdomen (while *Paleospinosus* has only one MP, *Latinozoros* has MPs on both T10 and T11) and the number of antennomeres (eight in *Paleospinosus*, nine in *Latinozoros*).

##### Systematic placement

Genus *Paleospinosus* was synonymized with *Octozoros* Engel, 2003 (that time a subgenus of *Zorotypus*) by Engel [24]. *Paleospinosus* is removed from that synonymy and reinstated as a valid genus in Spiralizoridae: Latinozorinae based on the arrangement of metatibial spurs.

##### Species included

*Paleospinosus hudae* Kaddumi, 2005.

##### Distribution

Jordan, Zarqa river basin (Lower Cretaceous, Albian).

#### Paleospinosus hudae Kaddumi, 2005

*Palaeospinosus hudae* Kaddumi, 2007: 218.

##### Comments

This species is known only from the holotype, which is an alate male [17]. The specimen is well preserved and allows us to observe important diagnostic characters, although the wing venation is just partly recognizable. Kaddumi [17] characterized venation as reduced, but Engel [24] commented it as a typical zorapteran venation. However, the presence or absence of forewing CuA_1_ is not recognizable for a the closer evaluation of the relationships of *Paleospinosus* with *Octozoros* and *Cretozoros*. Genitalia are not observable.

#### Mesozoic Zoraptera *incertae sedis*

##### *Zorotypus cretatus* Engel & Grimaldi, 2002

*Zorotypus cretatus* Engel & Grimaldi, 2002: 4.

###### Distribution

Myanmar, Kachin State, Myitkyina District, Hukawng Valley, Burmese amber (Upper Cretaceous, lower Cenomanian).

###### Comments

This species is known only from the apterous type male [21]. *Zorotypus cretatus* could not be classified due to poor fossil preservation and not observable diagnostic characters.

#### *Zorotypus dilaticeps* Yin, Cai, Huang & Engel, 2018

*Zorotypus dilaticeps* Yin, Cai, Huang & Engel, 2018: 127.

##### Distribution

Myanmar, Kachin State, Myitkyina District, Hukawng Valley, Burmese amber (Upper Cretaceous, lower Cenomanian).

##### Comments

*Zorotypus dilaticeps* is known only from an apterous (dealate) female type specimen [20]. It is a large species (3.9 mm), which can be distinguished from all other extinct and recent Zoraptera by its distinctive head morphology and the spination of the metafemur and metatibia. The inner margin of the metatibia is armed by six acute spines and seven spine-like setae. Such spination is unique among fossil Zoraptera and does not allow the classification to any extant genus; anyway, such arrangement does not need to be necessary for a genus-diagnostic character. Within recent groups of Zoraptera, a similar example of the secondary spination of metatibia appears in *Brazilozoros huxleyi* (Bolívar y Pieltain & Coronado, 1963), although other representatives of the genus do not have any spination [2, 13, 25]. Because the holotype specimen is a dealate female, placement of this taxon to a higher classification is not possible at the moment.

#### Identification key to fossil genera of Zoraptera (males only)

1. (6) Antennae composed of eight antennomeres; metatibia with two robust spurs, additional tiny apical spine could be developed on the inside of ventral surface; genitalia symmetrical.
2. (3) Ventral margin of metafemur with a deep longitudinal furrow; distal edge of S8 broadly emarginate with a peg-like projection in the middle …………………………………………… *Paleospinosus* Kaddumi, 2005, stat. restit.
3. (2) Ventral margin of metafemur entire or with only shallow longitudinal furrow; distal edge of S8 entire.
4. (5) Fore wing with CuA_1_ absent; ctenidium on T10 missing, mating hooks on both T10 and T11, genitalia with laterally developed rod-shaped accessory sclerites …………………………………………… *Cretozoros* gen. nov.
5. (4) Fore wing with CuA_1_ developed; ctenidium on T10 present; mating hook only on T11, genitalia without laterally developed rod-shaped accessory sclerites …………………………………………… *Octozoros* Engel, 2003, stat. nov.
6. (1) Antennae composed of nine antennomeres; metatibia with three robust spurs, additional apical spine on the inside of ventral surface not developed; genitalia asymmetrical
7. (8) Conical mating hooks on T10 and T11 *Burmazoros* gen. nov.
8. (7) Long procurved mating hook only on T10 *enozorotypus* Engel & Grimaldi, 2002

## Discussion

This study presents the first critical review of the so far described fossil species of Zoraptera described to date from the Mesozoic Era, which resulted in their classification based on the characters for which their diagnostic character was identified in recent species [13]. Currently, 11 Mesozoic Zoraptera species are recognized as valid (Tab. 2). Based on our morphological examination, nine species can be classified into five genera, i.e. *Burmazoros* gen. nov., *Cretozoros* gen. nov., *Octozoros* stat. nov., *Paleospinosus* Kaddumi, 2005, stat. restit. and *Xenozorotypus* Engel & Grimaldi, 2002. The current Zoraptera system is based on the characters of male genitalia (Additional file 1: Figure S1). The principal diagnostic character is the symmetry of the male copulatory organ, with representatives of Zorotypidae having asymmetrical genitalia and without a developed basal plate, while Spiralizoridae has symmetrical genitalia with a developed basal plate. The division into subfamilies also reflects the basic structural plan of the genitalia, with the number and degree of symmetry of the sclerites (Zorotypinae vs. Spermozorinae) and the development of the intromittent organ [13]. Higher taxonomic units are defined on these characters (see the identification key above). Male genitalia have not yet been properly studied and described in fossil Zoraptera, and can be observed only on two specimens, each belonging to a different genus. In *Cretozoros pusillus*, the everted genitalia are difficult to homologize [16]. In *Burmazoros denticulatus*, genitalia are partly visible in the holotype; however, they were not described by the authors. Within the extensively studied material of Mesozoic Zoraptera, we managed to find three male specimens with partially observable genitalia, which enabled homologizing their genitalia with those of the recent groups and, subsequently, the inclusion of most Mesozoic species into the current classification of the order.

Within the framework of recent phylogenetic studies [2, 6, 13], it was possible to homologize phylogenetically supporting characters with several external morphological characters, which, according to the current knowledge, also bear a phylogenetic signal. The most important are the characters on the hind legs, which in Zoraptera occur in three states as follows: three spurs, two spurs, or no spur (the latter with only small apical bristles which may be completely absent [8, 13], see Additional file 1: Figure S1). In recent groups, the number of spurs correlates with the character of male genitalia within molecularly supported monophyletic groups [13]. In the genera *Octozoros*, *Burmazoros* and *Cretozoros*, the number of spurs on the hind legs fully correlates with the morphology of male genitalia, as is documented in the current system constructed using the molecular phylogenetic approach, based on which we can also classify species with not observable genitalia. Three newly defined (or reinstated) genera *Octozoros*, *Cretozoros*, and *Paleospinosus* include individuals with two spurs on hind legs, and since the character of their genitalia correlates with the subfamily Latinozorinae, we classify them in this group (Tab. 1, 2, Fig. 4). In newly erected *Burmazoros*, the character of asymmetrical genitalia correlates with three spurs in metalegs as in the recent Zorotypinae, therefore we classify it in this group. The same applies to *Xenozorotypus burmiticus*, which is provisionally classified in the same subfamily based on the character of metatibie, although the morphology of the male copulatory organ is not known. The systematic placement of this taxon can be later refined when we find individuals with at least partially visible genitalia.

**Table 1.**
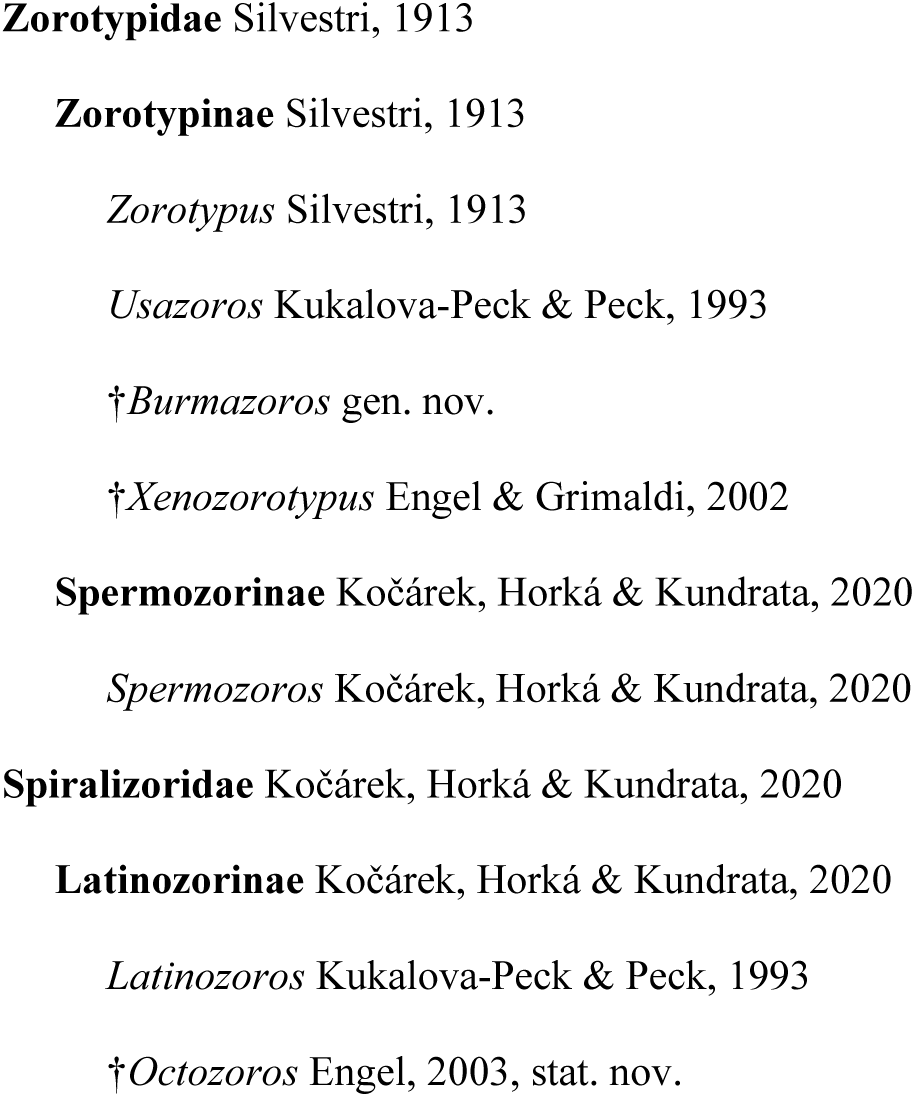
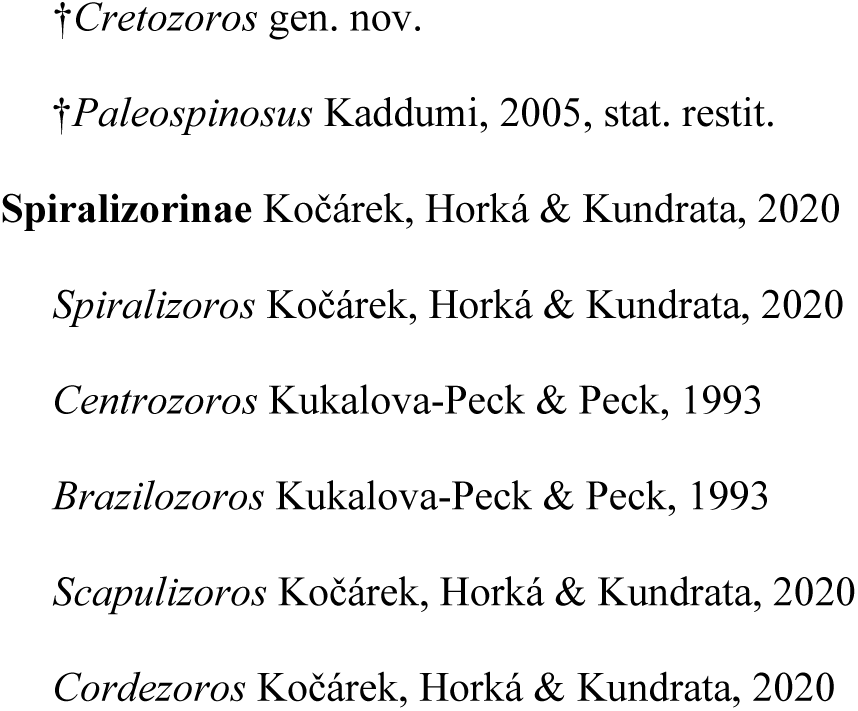
An updated higher-level classification of Zoraptera.

**Table 2.**
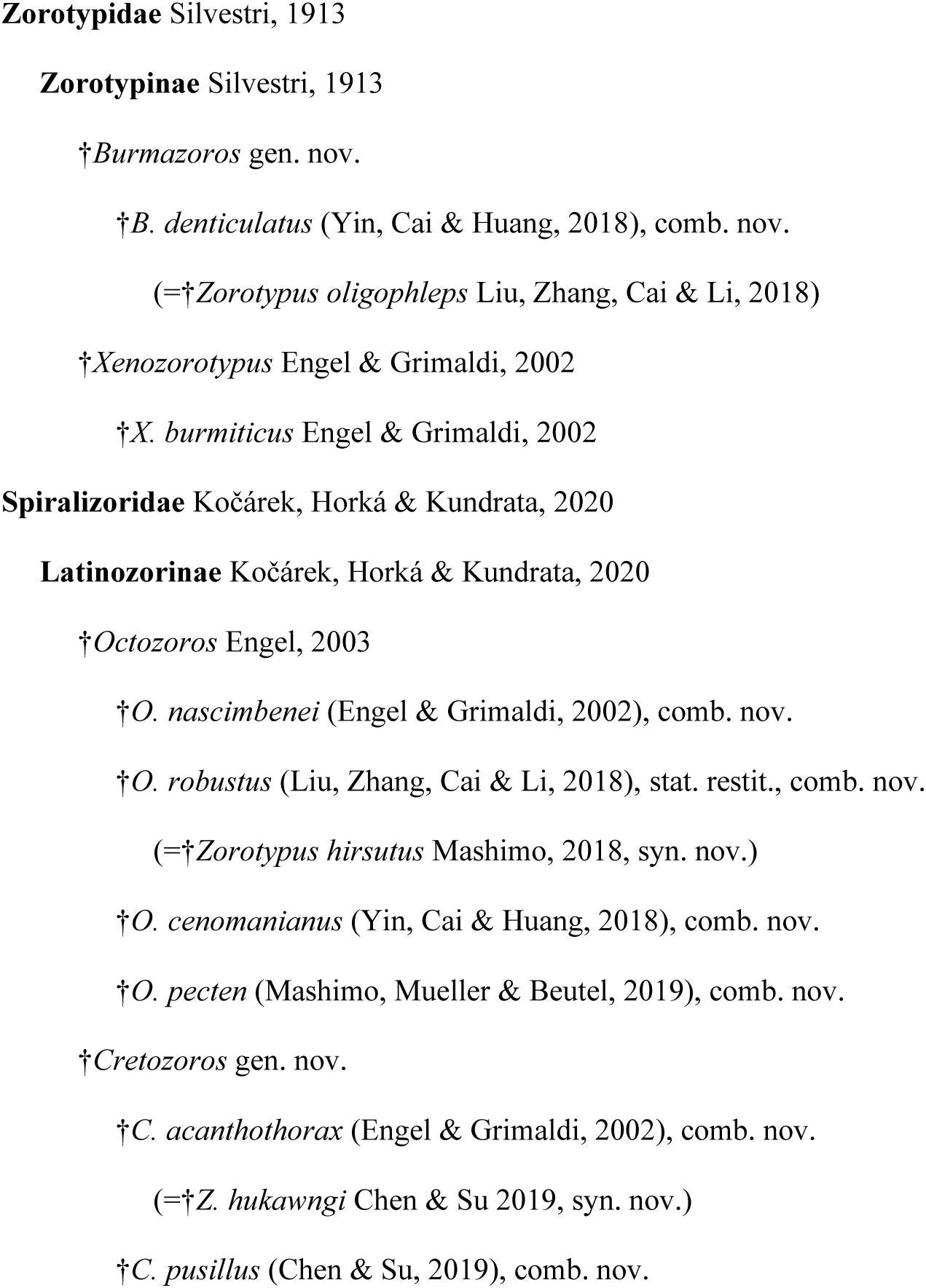
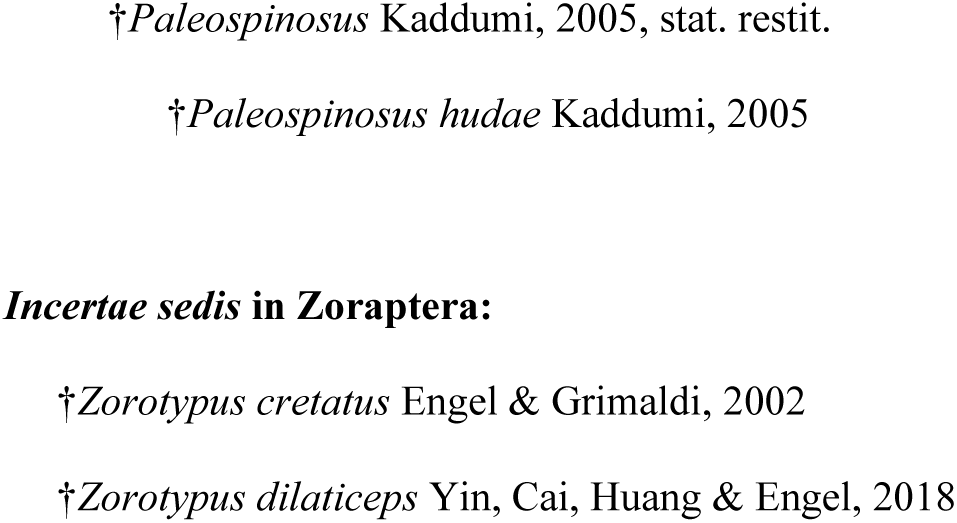
The checklist of Mesozoic Zoraptera.

The phylogenetic significance of wing venation remains questionable. Kukalova-Peck & Peck [26] proposed the first generic classification of recent Zoraptera based on wing venation, but this was not accepted by the community [21], and subsequent authors due to both high interspecific similarity and intraspecific variability of venation. Since then, no author has evaluated the importance of wing venation for phylogenetic reconstruction. A new analysis in light of recently established molecular-phylogenetic relationships is highly desirable. One of the few apomorphies, already defined by Kukalova-Peck & Peck [26], is the absence of CuA_1_ vein in the genus *Latinozoros*, and it seems to indeed bear a phylogenetic signal. The absence of this vein is associated with recent representatives of the genus *Latinozoros* (Kočárek & Horká [6] and unpublished authors’ observations), and its absence was also found in the here-defined genus *Cretozoros*. Therefore, we assume its close relationship with the recent genus *Latinozoros* (Fig. 4), which is supported by the morphologically similar type of male genitalia. This vein is already present in the genus *Octozoros*, and since the male genitalia also show a different character, we consider this group to be sister to the clade *Latinozoros*+*Cretozoros* (Fig. 4). Regarding the fossil genus *Xenozorotypus*, we tentatively classify it in Zorotypidae: Zorotypinae based on the number of spurs on the hind legs. It has a unique arrangement of the veins on the hind wing (presence of M_3+4_ vein). Characteristically reduced venation is present in *Burmazoros* gen. nov., which is classified here in Zorotypidae: Zorotypinae. It has fore wings with missing CuA, and hind wings with missing M_1+2_ and M fused with R [19, 22].

For some recent genera, which seems also valid for some fossil genera, a diagnostic character is the presence of median projections (MPs) in males on the posterior parts of abdominal tergites 10 and 11 [13]. There are several states known for this character in fossil Zoraptera, i.e., the presence of MP on both T10 and T11, the presence of only T11, and the absence of both MPs. In recent groups, this character is constant for all species of some higher taxonomic groups (e.g., all recent Zorotypinae have developed MP T11 only, and recent species of Latinozorinae and Spermozorinae have both MP T10 and T11). However, in recent representatives of the genus *Zorotypus*, we encounter MP T10+T11, MP T11, and also the complete absence of both MPs. The importance of this character for the classification thus remains questionable and is only relevant for some groups. Among fossil species, we also encounter the variability of this character. Within the fossil Latinozorinae, the representatives of *Octozoros* have only MP11 developed, and in the genus *Cretozoros* gen. nov., both MP10 and MP11 are developed, although in some species of this genus, the detection of MP10 is uncertain and it may be absent. *Paleospinosus* has developed only one MP, but it is not clear from the fossil whether it is the projection of T10 or T11. *Burmazoros* (Zorotypidae: Zorotypinae) has a small conical MP on both T10 and T11; on the other hand, *Xenozorotypus*, classified in the same subfamily, has only a T10 projection. Recent *Zorotypus* spp. have a mating hook on T10 or T11, or on both T10 and T11 or fused T10+T11 [4, 8, 13].

Based on current knowledge, we can include *Cretozoros*, *Burmazoros*, *Octozoros*, and *Paleospinosus* in the zorapteran phylogenetic system with a high degree of certainty (Fig. 4). Besides that, we place the genus *Xenozorotypus* provisionally in Zorotypidae: However, in Zorotypinae the systematic placement of this taxon will need to be confirmed in the future (Fig. 4). We provisionally keep the remaining two described Mesozoic species in the genus *Zorotypus* (*Z. cretatus*; *Z. dilaticeps* as *incertae sedis*, Tab. 2). For their proper classification, it will be necessary to wait until we find further individuals with well-visible diagnostic morphological structures. We consider the morphology of male genitalia to be the most important in this case.

An important aspect of the study of fossils in amber, which does not only concern Zoraptera, is the need for very careful morphological comparison of examined material with already described species, as very often the newly described species are only additional specimens of previously described species. The reasons are the different levels of preservation of the fossils, the different positions of the fossils, and the resulting different levels of observability of the structures, but also secondary changes in shape caused by compressions [27]. A significant aspect is also the usual observation of only one sex, which may have some different sex-specific characteristics. In the case of Zoraptera, the situation is additionally complicated by winged vs. wingless specimens and by the thin cuticle of their bodies, which is easily deformed in the amber. At the species level, characters related to the pattern of distribution or size of the spurs on the hind legs are useful. Above all, the metafemur spur distribution pattern seems to be a good species-diagnostic character. However, its usefulness for higher classification is minimal, as shown in the phylogenetic study of recent Zoraptera [13]. We highly recommend sticking to the approach that if the examined individuals have the same pattern of spines on the metafemur and spines on the metatibia (position of the spurs and their length) as the compared species, and if the specimen does not show any other prominent apomorphy (e.g., character of wing venation, spines on the chest, etc.) we consider them the same species. Caution is especially necessary when assessing the shape and length of cerci and antennomeres, which are both very susceptible to deformations and can look different even in specimens of the same species. To correctly assess these characters, it is recommended to have several specimens available for comparison.

## Material and methods

We critically reviewed all relevant literature on fossil Mesozoic Zoraptera to assess their systematic placement. In addition to the already published information, we studied in detail several newly reported specimens of the most critical taxa. The studied material is preserved in fossil resin originally produced by representatives of the tree family Araucariaceae [28]. Studied amber pieces were found in the surroundings of Tanai Village (26°21’N, 96°43’E) in the Hukawng Valley of Myanmar [29–31], but precise mining sites are unknown due to mixing of samples obtained from local miners. Deposits in Tanai Village have been investigated and dated in detail by Cruickshank & Ko [30] and Shi et al. [32]; the age has been estimated at approximately 99 Ma (98.8 ± 0.6; Lower Cenomanian) based on U-Pb dating of zircons from the volcaniclastic matrix of the amber [32]. The studied specimens have been collected before 2017 and legally exported from Myanmar (see the discussion in Haug et al. [33]). The amber pieces containing the specimens were ground, polished, and then examined with a Leica Z16 APO macroscope (Leica Microsystems, Wetzlar, Germany) equipped with a Canon 6D Mark II camera (Canon Inc., Tokyo, Japan). Micrographs of 20 to 30 focal layers of the same sample were combined with Helicon Focus software (Helicon Soft Ltd., Kharkiv, Ukraine) and finally processed with Adobe Photoshop CS6 Extended v13 (Adobe Inc., San Jose, California).

In this study, the following fossil samples deposited at the Department of Biology of the University of Ostrava, Czech Republic were examined: *Octozoros cenomanianus* (Yin, Cai & Huang, 2018), comb. nov.: PK344Bu (alate male); *O. robustus* (Liu, Zhang, Cai & Li, 2018), comb. nov.: PK348Bu (alate female), PK343Bu (apterous male), PK336Bu (apterous female), PK164Bu (alate female); *Cretozoros acanthothorax* (Engel & Grimaldi, 2002), comb. nov.: PK342Bu (alate male), PK347Bu (alate male), PK340Bu (alate male), PK346Bu (alate female). Illustrations of subfamily diagnostic characters were adopted from Kočárek et al. [13] and supplemented by the microphotograph of hind leg of *Spermozoros weiweii* (Wang, Li & Cai, 2016), collected in Brunei Darussalam (Ulu Temburong NP, Sungai Apan II, N 4°33.22950’ E 115°10.59277’, 12.–19.ii.2015, P. Kočárek & I. Horká leg). The higher classification and morphological terminology of Zoraptera follow Kočárek et al. [13]. Geological periods and epochs follow the International Chronostratigraphic Chart v2023/09 [34].

Abbreviations used in the text: Cu, cubitus vein; CuA, anterior cubitus vein; M, media vein; MP, median projection; R, radius vein; S, abdominal sternite; T, abdominal tergite.

## Conclusions

Zoraptera, or angel insects, represent one of the less numerous insect orders, with only 47 extant and 16 extinct described species. Zoraptera are small, soft-bodied, primarily winged polyneopteran insects and their uniformity in general morphology has led to the persistence of a conservative classification of extant Zoraptera, with only a single nominotypical genus *Zorotypus* Silvestri, 1913 in a single family Zorotypidae for more than a century. Kočárek et al. [13] and Matsumura et al. [8] conducted molecular phylogenetic studies using a combination of nuclear and mitochondrial markers. Both of these independent analyses revealed two major phylogenetic lineages, which Kočárek et al. [13] classified as families Zorotypidae and Spiralizoridae, each of them being further subdivided into two robustly supported subfamilies. The current system of extant Zoraptera is based on the results of molecular phylogeny combined with the morphology of male genitalia, and supplemented by the characters on the male abdomen and a number of metatibial spurs. However, fossil representatives of Zoraptera have not yet been classified into the modern system and most of them remained in the collecting genus *Zorotypus* Silvestri, 1913, because the genitalia were not observable or examined in detail. Unfortunately, molecular-genetic methods cannot be applied to insect fossils trapped in amber; therefore, their systematic placement must be inferred from the morphological characters.

Zoraptera represent an old evolutionary lineage with a Paleozoic origin, and more lines of evidence suggest that Zoraptera were already diversified and widely distributed in Gondwana at the time of the break-up of this supercontinent. Together, 12 fossil species of Zoraptera are currently recognised from Mesozoic. One species is reported from the lower Cretaceous Jordanian amber, and 11 species are known from the upper Cretaceous amber of northern Myanmar. Fossil zorapterans are classified partly in *Zorotypus sensu stricto*, partly in the monotypic genus *Xenozorotypus* Engel & Grimaldi, 2002, and partly in the exclusively fossil subgenus *Octozoros* Engel, 2003, which was erected for species with eight antennomeres. All of these taxa were included in a single family Zorotypidae.

In this study, for the first time, we described and critically evaluated male genitalia and other main diagnostic characters of all available Mesozoic Zoraptera. Our results lead to the first proposal of the generic classification of Mesozoic Zoraptera. We described two new genera, *Cretozoros* gen. nov. and *Burmazoros* gen. nov., reinstated *Paleospinosus* Kaddumi, 2005, stat. restit. from synonymy with the subgenus *Octozoros* Engel, 2003 (in Zorotypus), and elevated *Octozoros* Engel, 2003 to a genus level. *Cretozoros* gen. nov., *Paleospinosus* Kaddumi, 2005, stat. restit., and *Octozoros* stat. nov. were classified in Spiralizoridae:

Latinozorinae, while *Burmazoros* gen. nov. and *Xenoburmiticus* Engel & Grimaldi, 2002 were classified in Zorotypidae: Zorotypinae. Altogether, it was possible to classify nine of the 11 currently recognized species of Mesozoic Zoraptera. *Zorotypus hukawngi* Chen & Su, 2019 was synonymized with *Cretozoros acanthothorax* (Engel & Grimaldi, 2002), comb. nov., and *Zorotypus hirsutus* Mashimo, 2018 was synonymized with *Octozoros robustus* (Liu, Zhang, Cai & Li, 2018), stat. restit., comb. nov., which was simultaneously restituted from the synonymy with *Octozoros cenomanianus* (Yin, Cai & Huang, 2018), comb. nov.

The classification of Mesozoic Zoraptera into the modern system enables us to better understand the diversity of their internal lineages during the early evolution of this enigmatic insect order.

## Acknowledgments

This research was supported by project GACR 22-05024S (Evolution of angel insects (Zoraptera): from fossils and comparative morphology to cytogenetics and transcriptomes). The authors also thank the staff of the Kuala Belalong Field Studies Centre for their support during the stay of our research team at the Centre and we thank the Universiti Brunei Darussalam for permission to collect Zoraptera.

## Authors’ contributions

P.K. conceived the study, processed the fossils and microphotographs, P.K. and I.K. prepared and composed illustrations, P.K. wrote the paper with contributions from I.K. and R.K.

## Funding

The research that led to these results received funding from the Czech Science Foundation under Grant Agreement No22-05024S.

## Data availability statement

All data generated or analysed during this study are included in this published article.

## Declarations

### Ethics approval and consent to participate

No applicable.

### Consent for publication

No applicable.

### Competing interests

The authors declare that they have no conflicts of interest.

## Supplementary information

Supplementary File S1: Identification keys to Zoraptera families and genera of Mesozoic Zoraptera

Supplementary Figure S1. Comparison of diagnostic characters for currently recognized subfamilies.

## Notes

### Competing Interest Statement

The authors have declared no competing interest.

